# Development and cross-cohort evaluation of a multimodal deep-learning model for ER, PR and HER2 status classification in breast cancer

**DOI:** 10.64898/2026.11.01.761587

**Authors:** Binh Minh Tran, Van An Nguyen, Quynh Anh Dang, Ngoc Minh Hai Nguyen, Quang Anh Nguyen, Hai Nam Tran, Dang Hung Tran, Thi Bich Van Nguyen

## Abstract

**Background:** Estrogen receptor (ER), progesterone receptor (PR) and HER2 status guide systemic treatment in breast cancer. Multimodal models are rarely benchmarked against key biomarker molecular baselines or evaluated for cross-cohort calibration. We assessed incremental value, dependence on label-proximal HER2 transcripts, and transport of discrimination and calibration.

**Methods:** A model combining whole-slide images, transcriptomes, age and stage was developed in TCGA-BRCA (1051 patients; ER/PR/HER2 analysis sets: 1001/998/1006) and evaluated in CPTAC-BRCA (120 patients; 118/113/95). CPTAC had been observed during earlier development and included 13 HER2-positive cases. Analyses included paired model comparisons, transcript removal, alternative loss functions and recalibration. Internal discrimination used pooled out-of-fold predictions; external discrimination used fold-mean estimates, with ensemble calibration reported separately. Internal calibration and macro-AUC uncertainty were interpreted as exploratory.

**Results:** Internal AUCs were 0.943 for ER, 0.901 for PR, and 0.860 for HER2; external AUCs were 0.952, 0.937, and 0.918, respectively. External macro-AUC was 0.936 for the multimodal model and 0.957 for logistic regression using ESR1, PGR, and ERBB2 (difference −0.021). Removal of the extended 29-transcript 17q12–q21 region changed HER2 AUC by −0.035 [−0.058 to −0.013] (95% CI) internally and by −0.042 [−0.089 to −0.007] externally relative to the base-line multimodal model. For external 15-model ensemble predictions, recalibrated HER2 expected calibration error (ECE) was 0.236, exceeding the ECE reference ceiling of 0.125, while calibration-in-the-large was −1.657 [−2.373 to −1.023] (profile-likelihood 95% CI), indicating systematic overprediction. ER and PR also showed overprediction. In paired fold-mean comparisons, HER2 ECE and Brier score favoured the three-transcript model (ESR1, PGR, and ERBB2).

**Conclusions:** A multimodal deep-learning model achieved high discrimination across cohorts but did not demonstrate clear incremental value over a par-simonious three-transcript baseline, while cross-cohort calibration remained a substantial challenge. These findings highlight the importance of evaluating incremental value, calibration, and clinical utility alongside discrimination when assessing AI-based prediction models for medical use.

## 1 Background

Breast cancer is the most commonly diagnosed malignancy among women worldwide, with an estimated 2,300,000 new cases in 2022 [1]. Estrogen receptor (ER), progesterone receptor (PR), and human epidermal growth factor receptor 2 (HER2) status guide the selection of endocrine therapy, HER2-targeted therapy, and cyto-toxic treatment [2, 3]. In routine practice, receptor status is assessed primarily by immunohistochemistry (IHC), with in situ hybridisation (ISH) used to resolve equivocal HER2 results. Although IHC/ISH remains the reference standard, pre-analytical variation, staining quality, assay standardisation, and interpretation can affect results, particularly at low levels of HER2 expression [2–5].

Information associated with receptor status is reflected across multiple biological scales. Histomorphological patterns in haematoxylin and eosin (H&E) whole-slide images contain features associated with ER, PR, and HER2 status, and deep-learning models have demonstrated the ability to extract these signals from routine pathology images [6–11]. Transcriptomic data provide a complementary molecular representation, with label-proximal transcripts such as ESR1, PGR, and ERBB2 closely associated with the corresponding phenotypes [12–15]. For HER2 positivity, this signal may extend beyond ERBB2 to neighbouring genes within the co-amplified 17q12– q21 region [16–18]. These complementary image-based and transcript-based signals provide a rationale for multimodal modelling [19, 20].

Artificial intelligence models that integrate histology, transcriptomics, and clinical information may offer additional support for receptor assessment. One potential application is laboratory quality assurance. In settings where digital pathology and tumour RNA sequencing are already available, model-based estimates could provide an orthogonal check to flag potentially discordant receptor results for further review before clinical reporting. Such a system would not replace IHC/ISH, but could help identify cases requiring closer inspection. This application remains hypothetical, however, and requires evidence that a multimodal model adds value beyond simpler molecular predictors and that its probability estimates remain reliable across patient cohorts.

Three related evidence questions therefore arise. First, complex multimodal models should be benchmarked against strong, biologically informed, parsimonious molecular baselines because greater modelling complexity does not necessarily improve clinical prediction performance [21]. Second, high HER2 discrimination may partly reflect reliance on label-proximal transcripts within the 17q12–q21 amplicon, making it important to examine how performance changes when these signals are removed [16–18]. Third, high discrimination does not ensure well-calibrated probabilities, and calibration and decision-curve analysis provide complementary information on probability accuracy and potential clinical usefulness [22, 23].

Accordingly, this study had three objectives:

(1) to develop and cross-cohort evaluate a multimodal model combining whole-slide images, tumour transcriptomes, age, and stage, and to quantify its incremental predictive value over parsimonious molecular and component baselines;
(2) to assess the model’s dependence on label-proximal molecular information, particularly the 17q12–q21 region for HER2 prediction, using transcript-removal and transcript-restriction experiments; and
(3) to assess the cross-cohort transport of predicted probabilities from TCGA-BRCA to CPTAC-BRCA, with emphasis on calibration and exploratory decision-curve performance. Reporting was guided by TRIPOD+AI [24].

## 2 Methods

### 2.1 Study design, data sources and reporting

This retrospective multi-cohort study developed a multimodal prediction model for ER, PR and HER2 status in TCGA-BRCA using patient-level cross-validation and evaluated it in CPTAC-BRCA without model refitting. Model performance on CPTAC-BRCA had been examined during an earlier development cycle; the external evaluation was therefore not fully independent, as detailed in Section 2.8.

The final analyses addressed three study questions and included calibration assessment and decision-curve analysis. The final reanalysis comprised 208 runs, with configuration choices recorded before execution; prespecification refers to this reanalysis rather than the earlier development cycle. Reporting follows TRIPOD+AI [24]. The reporting checklist (Additional file 1) and PROBAST+AI self-assessment [25] (Additional file 2) are provided.

Only de-identified data from public repositories were analysed. The study involved no participant contact, intervention or new specimen collection.

### 2.2 Data sources and participants

Development data comprised clinical records, RNA sequencing (STAR gene counts) and diagnostic H&E whole-slide images from TCGA-BRCA [26]. External evaluation used the corresponding modalities from the CPTAC-BRCA cohort described by Krug et al. [27]. The study design and evaluation workflow are shown in Figure 1.

**Fig 1.**
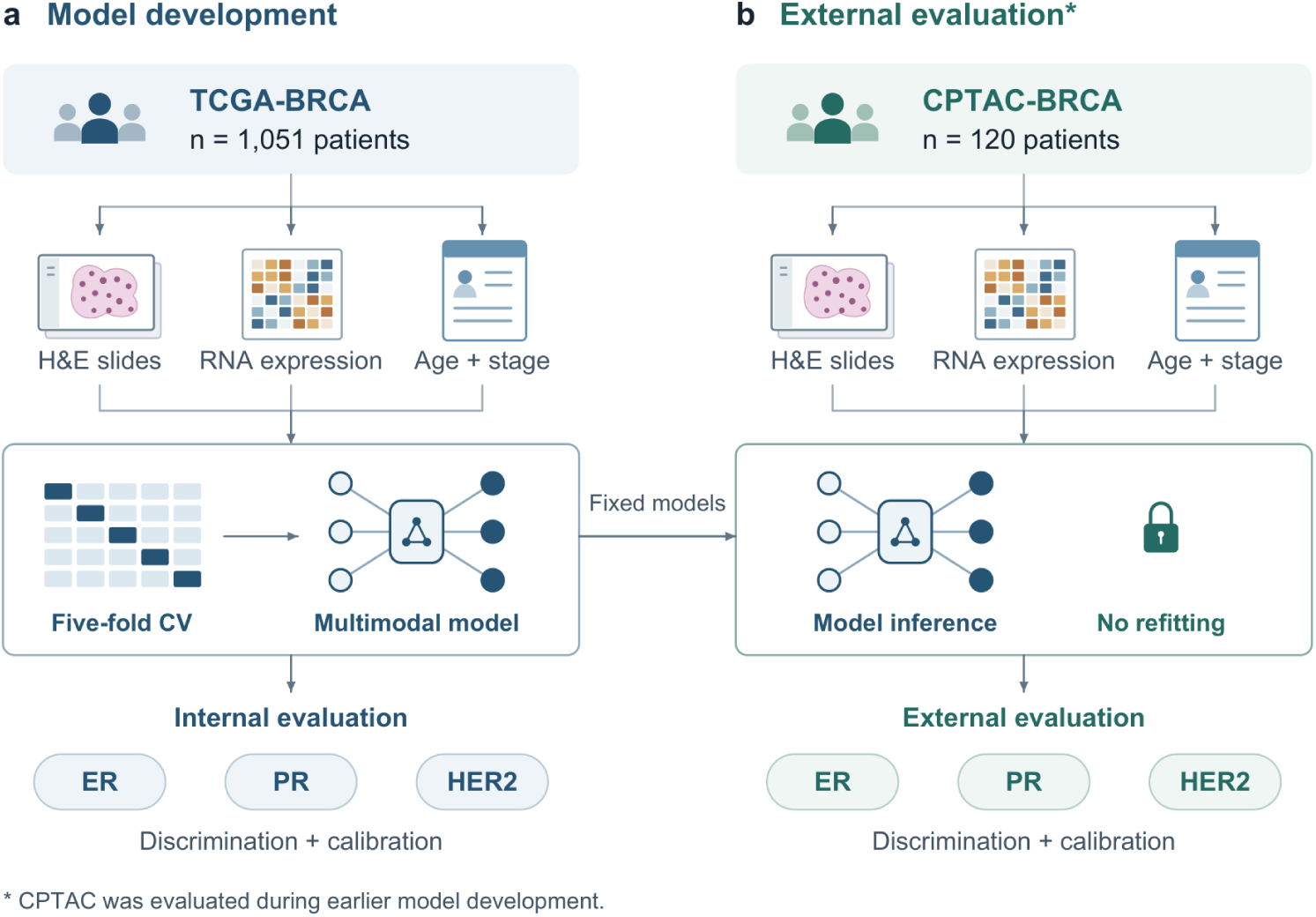
Study design. Model development and internal evaluation in TCGA-BRCA, followed by external evaluation in CPTAC-BRCA without model refitting. CPTAC-BRCA had been evaluated during an earlier development cycle.

The final TCGA-BRCA development cohort included 1051 patients with primary tumours and matched data from all three modalities. Three of the 1054 patients with diagnostic whole-slide images were excluded because matched RNA sequencing or clinical data were unavailable. For 66 patients with multiple diagnostic slides, tiles were pooled into a single patient-level bag before data splitting.

In CPTAC-BRCA, 122 patients had RNA sequencing and clinical data. Precomputed whole-slide feature bags were unavailable for two patients, leaving 120 for external evaluation.

Unobserved receptor labels were masked rather than imputed. Patients remained eligible for each endpoint with an observed label; endpoint-specific sample sizes therefore reflect label availability rather than separate patient cohorts. In TCGA-BRCA, labels were unobserved for 50 ER, 53 PR and 45 HER2 cases, yielding analysis sets of 1001, 998 and 1006 patients, respectively. In CPTAC-BRCA, the corresponding numbers of unobserved labels were 2, 7 and 25, yielding analysis sets of 118, 113 and 95 patients.

### 2.3 Outcome definition and reference standard

The outcomes were binary ER, PR and HER2 status, derived from the available clinical and pathology annotations in each cohort. In clinical practice, ER and PR are assessed by immunohistochemistry (IHC), whereas HER2 assessment uses IHC and in situ hybridisation (ISH), with ISH used to resolve equivocal HER2 IHC results [2, 3]. HER2 label provenance differed between cohorts. The primary TCGA-BRCA label rule (L0) applied an ordered hierarchy: definitive IHC scores (0/1+ negative, 317 patients; 3+ positive, 88), ISH records (270), unanimous GDC ERBB2 ISH/FISH records (2), curated IHC annotations (239), and the 2012 TCGA HER2 status field (90). Each source resolved only cases left unresolved by preceding sources; equivocal values were insufficient, and earlier sources prevailed when records disagreed. L0 yielded 1006 labelled patients, including 184 positive cases, and 45 unresolved patients, whose HER2 labels were treated as unobserved.

In CPTAC-BRCA, HER2 labels were obtained from a single curated clinical field. Eight patients had equivocal IHC 2+ results without confirmatory ISH. Their HER2 labels were treated as unobserved and masked for the HER2 endpoint.

Alternative HER2 label rules were examined in sensitivity analyses: L1 prioritised available ISH/FISH results over IHC-based labels, whereas L2 retained only high-confidence labels from definitive IHC (0/1+ or 3+) or ISH/FISH and treated all other cases as unobserved. All models remained trained on L0; these analyses reassessed performance under alternative label definitions rather than retraining the models.

### 2.4 Predictors

#### 2.4.1 Clinical variables

Clinical predictors comprised age at diagnosis and AJCC stage, the two clinical fields with a shared schema across cohorts. Missing age values were imputed using the median estimated from the corresponding training-fold data. Stage was one-hot encoded, with missing values represented by an Unknown category. No receptor-status or molecular-subtype variables were included as clinical predictors.

#### 2.4.2 Transcriptomic profile

Count preprocessing comprised aggregation of Ensembl identifiers to gene symbols and conversion to log2(CPM + 1). Expression matrices were aligned to 20,344 gene symbols shared between TCGA-BRCA and CPTAC-BRCA. These shared symbols formed the candidate pool. Within each TCGA inner-training fold, the 5000 genes with the highest variance were ranked by mutual information separately for ER, PR and HER2 using observed labels; predefined withheld genes were removed before variance filtering. The union of the top 400 genes per outcome formed the full selected transcriptomic panel used by G0 and R5b, with a mean size of 832 genes (range, 820–844). These TCGA-selected predictors were retained for external evaluation. This selected panel differed from the 5000 highest-variance genes used by the elastic-net comparator R5.

In TCGA-BRCA, a deterministic rank-based inverse-normal transformation was fitted on inner-training data and applied unchanged to inner-validation and heldout test data. To accommodate differences between expression platforms, ranks were recomputed within CPTAC-BRCA without using receptor labels. This transductive preprocessing step depended on the external cohort and does not directly support isolated single-patient inference. Fold-specific panels and deterministic preprocessing records were retained, as described in Section 2.11.

#### 2.4.3 Whole-slide images

Diagnostic whole-slide images were divided into non-overlapping 256×256-pixel tiles at an apparent magnification of 20×. Tile embeddings were extracted using the publicly released UNI2-h encoder (ViT-H/14; 1536-dimensional embeddings) [28, 29] through the Trident toolkit [30]. The encoder was used without fine-tuning, and no stain normalisation was applied. Patient-level bags contained at most 2048 tiles, with up to 512 tiles sampled from each bag per training or inference pass.

### 2.5 Model architecture

The model combined clinical, transcriptomic and whole-slide image (WSI) inputs through modality-specific encoders, self-attention fusion and a shared multitask prediction network (Figure 2).

**Fig 2.**
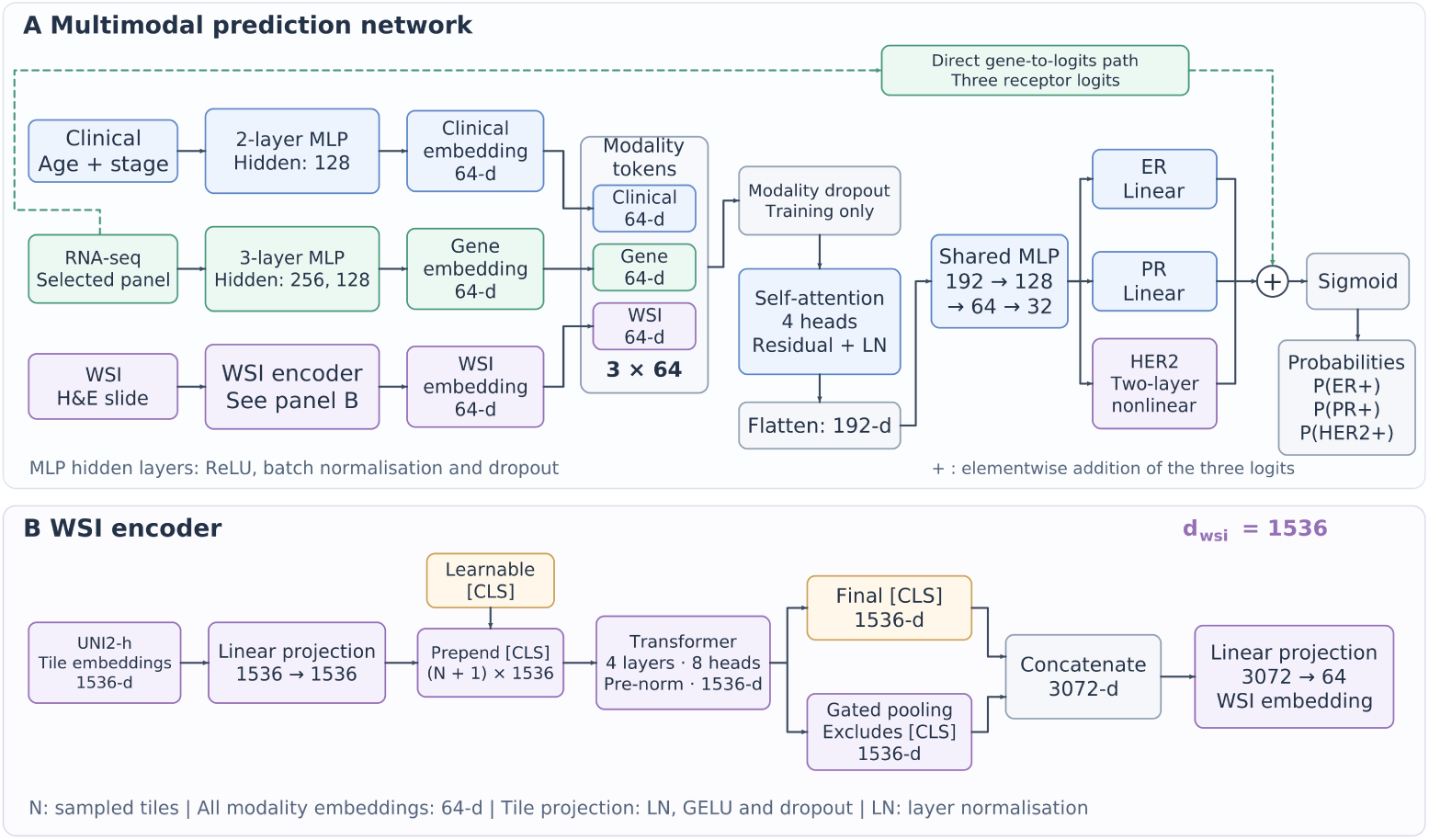
Multimodal multitask architecture for receptor status prediction. (A) Clinical, transcriptomic and WSI encoders produce 64-dimensional modality embeddings. Modality dropout is applied during training, followed by four-head self-attention with a residual connection and layer normalisation. The fused tokens are flattened into a 192-dimensional vector and passed through a shared MLP trunk and receptor-specific heads. The direct gene-to-logits path contributes to all three receptor logits before sigmoid conversion to probabilities. (B) The WSI encoder processes precomputed UNI2-h tile embeddings using a four-layer, eight-head Transformer with *d*_wsi_ = 1536. The final classification-token state and gated-attention pooled vector are concatenated (3072 dimensions) and projected to *d_f_* = 64. MLP, multilayer perceptron; WSI, whole-slide image; CLS, classification token; LN, layer normalisation; ER, estrogen receptor; PR, progesterone receptor; HER2, human epidermal growth factor receptor 2.

Clinical variables were encoded using a two-layer multilayer perceptron (MLP) with a hidden width of 128. Gene-expression profiles were encoded using a three-layer MLP with hidden widths of 256 and 128. Both branches produced embeddings of dimension *d*_f_ = 64. Each hidden layer was followed by ReLU activation, batch normalisation and dropout.

For the WSI branch, each patient was represented by a bag of precomputed 1536-dimensional UNI2-h tile embeddings, as described in the Predictors subsection. Tile embeddings were projected to dimension *d*_wsi_ = 1536 using a learned linear layer followed by layer normalisation, GELU activation and dropout. A learnable classification token ([CLS]) was prepended to the tile sequence, which was processed by a four-layer pre-norm Transformer encoder with eight attention heads [31]. Gated-attention pooling was applied to the resulting tile-token representations, excluding the classification token, to obtain an attention-weighted summary [32]. This 1536-dimensional summary was concatenated with the 1536-dimensional final classification-token state and projected from 3072 dimensions to *d*_f_ = 64, yielding a patient-level WSI representation.

The three modality embeddings were stacked into a token matrix *S_i_ ∈* R^3*×df*^ for patient i and fused using four-head self-attention with a residual connection and layer normalisation:

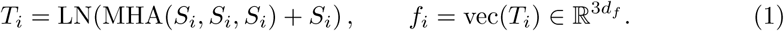

where MHA denotes multi-head attention, LN denotes layer normalisation, and vec denotes flattening of the attended modality tokens.

The 192-dimensional fused vector was passed through a shared MLP trunk with widths of 128, 64 and 32, with ReLU activation, batch normalisation and dropout after each layer. The resulting patient-level representation fed three receptor-specific prediction heads. ER and PR used linear heads, whereas HER2 used a two-layer head with an intermediate nonlinear activation. Fusion used modality dropout during training, and a direct gene-to-logits path complemented the fusion pathway. Final receptor logits were converted to probabilities using the sigmoid function.

### 2.6 Training objective

Focal loss for ER and PR [33] and asymmetric loss for HER2 [34] were retained from the initial development to address class imbalance and combined by homoscedastic uncertainty weighting [35]. Masking unobserved labels allowed a single multitask network to use all 1051 patients. An exploratory 2 × 2 comparison additionally included a binary-cross-entropy multimodal model and a three-transcript model fitted with focal/asymmetric objectives. Folds and preprocessing were shared across these configurations; factorial reporting was restricted to discrimination.

### 2.7 Training and internal validation

Internal evaluation used patient-level stratified five-fold cross-validation with three initialisations per fold and a 10% inner-validation subset for early stopping and threshold selection. Predictions were probability-averaged across the three initialisations within each fold, and internal metrics were calculated from pooled out-of-fold predictions. Configuration choices were fixed for the reanalysis but inherited from non-nested model selection; internal performance was therefore considered exploratory. Two additional five-fold partitions assessed stability under fixed configurations rather than repeated nested model selection. Partition and initialisation variability were summarised as descriptive ranges. Training–test AUC differences were reported descriptively and were not treated as estimates of selection-induced optimism.

### 2.8 External evaluation

Primary external discrimination was the mean of metrics calculated for five fold models, each averaging three initialisations. This fold-mean estimate differs from the internal pooled out-of-fold estimate. External discrimination and threshold-based performance were summarised across fold-specific models, whereas absolute calibration and decision-curve analyses used the 15-model ensemble to evaluate a single final probability per patient. For discrimination, the 15-model ensemble and one-initialisation-per-fold comparisons were sensitivity analyses. Weights were not refitted on CPTAC; thresholds and calibration mappings were estimated from TCGA. Label-free within-cohort ranking was the only CPTAC-dependent preprocessing. A TCGA-reference quantile-normalisation variant was assessed against a prespecified expression-unit comparability criterion.

CPTAC had been evaluated before the architecture was finalised in an earlier development cycle. Subsequent configuration selection used TCGA inner-validation data. The final results are therefore described as cross-cohort external evaluation rather than independent or confirmatory validation.

### 2.9 Comparators and transcript-removal experiments

Comparators comprised clinical-only and transcript-based logistic models, elastic-net regression [36], ridge regression on the full selected transcriptomic panel (the selected-panel ridge model, R5b), transcriptomic and WSI-only neural networks, and image– gene fusion models. Unless otherwise specified, the three-transcript model refers to logistic regression using ESR1, PGR and ERBB2 (R4). The full multimodal model is labelled R10 in the comparator analyses and G0 in the transcript-removal analyses. Per-patient predictions were retained for paired comparisons on common analysis sets. The primary contrast was external fold-mean macro-AUC for the multimodal model minus the three-transcript model. Transcript-removal experiments excluded predefined transcripts before feature selection; the restriction experiment retained only ESR1, PGR and ERBB2 in the transcriptomic branch. WSI and clinical branches were retained throughout [16–18]. The external HER2 AUC difference after removal of the extended 29-transcript 17q12–q21 region was the prespecified secondary contrast; other transcript-removal and restriction comparisons were exploratory.

Label-proximal transcripts were defined as ESR1, PGR and ERBB2 together with transcripts co-amplified or co-regulated with ERBB2 in the 17q12–q21 amplicon. The six-gene 17q12 core (G3) comprised ERBB2, GRB7, MIEN1, PGAP3, PNMT and STARD3. The extended 29-transcript 17q12–q21 region (G3b) is defined in Additional file 3 (Definitions sheet). The full predefined label-proximal set (G5) was the union of G3b and ESR1/PGR. Transcript-removal experiments excluded predefined subsets of these transcripts to assess dependence on label-proximal molecular information.

### 2.10 Statistical analysis

Discrimination was assessed using raw-probability AUC and area under the precision-recall curve (AUPRC). Maximum-F1 thresholds were selected from observed TCGA inner-validation probabilities. Thresholds were averaged across individual initialisations for internal assessment and selected from seed-averaged fold predictions for external assessment. A single resulting threshold per receptor was applied throughout each cohort without optimisation on CPTAC. Calibration measures comprised Brier score, expected calibration error (ECE), slope and calibration-in-the-large, the latter estimated with slope fixed at 1. Internal Platt recalibration [37] fitted each fold’s mapping to out-of-fold predictions from the other folds. For external ensemble analyses, a mapping fitted to all pooled TCGA out-of-fold predictions was applied after probability averaging. This pooled mapping also underpinned the reported external fold-mean head-to-head contrasts. Fold-specific inner-validation mappings were examined separately as sensitivity analyses. Internal recalibration and threshold-based performance were considered exploratory because the evaluation reused a common cross-validation partition.

ECE used quantile bins, with the number capped at the smaller of 10 and the event count. A finite-sample ECE reference ceiling was simulated from 5000 datasets matched on sample size and expected prevalence, using Beta-distributed probabilities and conditionally Bernoulli outcomes. Concentrations *κ* = 2, 5, 20 and 100 represented different probability dispersions; the largest scenario-specific 95th percentile was retained. This provides a reference for these scenarios, not a universal calibration test. Ceilings were calculated for pooled out-of-fold and external ensemble predictions, not fold-mean metrics. External absolute calibration and decision curves used ensemble predictions, whereas paired calibration contrasts explicitly identified their aggregation method.

HER2 provenance analyses compared high-confidence with other labels and ISH-derived with curated-IHC labels; disjoint strata were resampled independently. Decision-curve analysis [23] was exploratory, with no action-specific clinical thresh-old. Bootstrap intervals used 2000 patient-level replicates with shared indices between models within each receptor analysis set. Outcome-stratified resampling was used for discrimination, operating characteristics and free-fit calibration parameters; unstratified resampling was used for calibration-in-the-large, ECE, Brier score and net benefit. No replicate was discarded for the reported estimates. Macro-AUC replicates combined receptor-specific results by replicate index rather than a joint patient resample across outcomes; their intervals were interpreted descriptively. Alternative resampling schemes were examined where available. External calibration-in-the-large additionally used profile-likelihood intervals. Unless otherwise stated, bracketed intervals are 95% confidence intervals, reported as estimate [lower to upper]. Macro-AUC intervals remain descriptive, as specified above and in the relevant table legends. Performance estimates and intervals are reported to three decimal places, with additional precision retained where rounding would obscure a non-zero value or an interval boundary. Sample sizes were determined by available data, without an a priori calculation; limited event counts precluded stable subgroup analyses.

### 2.11 Validity and reproducibility

Programmatic checks assessed ten potential leakage pathways. A separate analysis evaluated the recoverability of TCGA submitting site from image features [38]. Fold-specific gene panels, predictions, deterministic preprocessing and internal replay records were retained. Analyses used Python 3.12.3, PyTorch 2.8.0 and scikit-learn 1.9.0 on an RTX PRO 4500 Blackwell GPU (32 GB). The project repository provides code, configuration records, gene panels and environment specifications [39]. The study archive is identified by DOI https://doi.org/10.5281/zenodo.22711622.

## 3 Results

### 3.1 Cohort characteristics

The development and external cohorts included 1051 and 120 patients, respectively (Table 1). Among patients with observed labels, ER, PR and HER2 positivity was 77.4%, 67.0% and 18.3% in TCGA and 68.6%, 60.2% and 13.7% in CPTAC. The internal HER2 set contained 1006 patients and 184 positive cases; the external set contained 95 patients and 13 positive cases.

**Table 1.** Cohort characteristics.

| **Characteristic** | **TCGA-BRCA (n = 1051)** | **CPTAC-BRCA (n = 120)** |
|---|---|---|
| **Demographics** |  |  |
| Age at diagnosis, years — median (IQR) | 59 (49–68) | 62 (51–71) |
| Age unknown, n (%) | 0 (0.0) | 15 (12.5) |
| Race — White, n (%) | 728 (69.3) | 77 (64.2) |
| Race — Black, n (%) | 167 (15.9) | 13 (10.8) |
| Race — Asian, n (%) | 61 (5.8) | 19 (15.8) |
| Race — other or not reported, n (%) | 95 (9.0) | 11 (9.2) |
| **Tumour stage** |  |  |
| Stage I, n (%) | 173 (16.5) | 4 (3.3) |
| Stage II, n (%) | 600 (57.1) | 69 (57.5) |
| Stage III, n (%) | 238 (22.6) | 32 (26.7) |
| Stage IV, n (%) | 19 (1.8) | 0 (0.0) |
| Stage unknown, n (%) | 21 (2.0) | 15 (12.5) |
| **Receptor status and analysis sets** |  |  |
| ER positive, n (% of labelled) | 775 (77.4) | 81 (68.6) |
| PR positive, n (% of labelled) | 669 (67.0) | 68 (60.2) |
| HER2 positive, n (% of labelled) | 184 (18.3) | 13 (13.7) |
| Analysable N (ER / PR / HER2) | 1001 / 998 / 1006 | 118 / 113 / 95 |
| **Events and precision of the analysis sets** |  |  |
| ER — events / non-events (AUC 95% CI width) | 775 / 226 (0.043) | 81 / 37 (0.079) |
| PR — events / non-events (AUC 95% CI width) | 669 / 329 (0.043) | 68 / 45 (0.083) |
| HER2<sup>a</sup> — events / non-events (AUC 95% CI width) | 184 / 822 (0.066) | 13 / 82 (0.166) |

Receptor positivity is calculated among patients with an observed label; other percentages use the full cohort. AUC 95% CI widths are those of Table 2A. IQR, interquartile range.
<sup>a</sup> External HER2 estimates are based on 13 positive cases and should be interpreted cautiously because precision is limited.

### 3.2 Discrimination and operating characteristics

Internal AUCs were 0.943, 0.901 and 0.860 for ER, PR and HER2; corresponding external fold-mean AUCs were 0.952, 0.937 and 0.918 (Table 2A). External AUPRCs were 0.976, 0.962 and 0.784. At thresholds selected using the development cohort, external fold-mean HER2 sensitivity was 0.923, specificity 0.578, positive predictive value 0.270 and negative predictive value 0.980 (Table 2C).

**Table 2.**
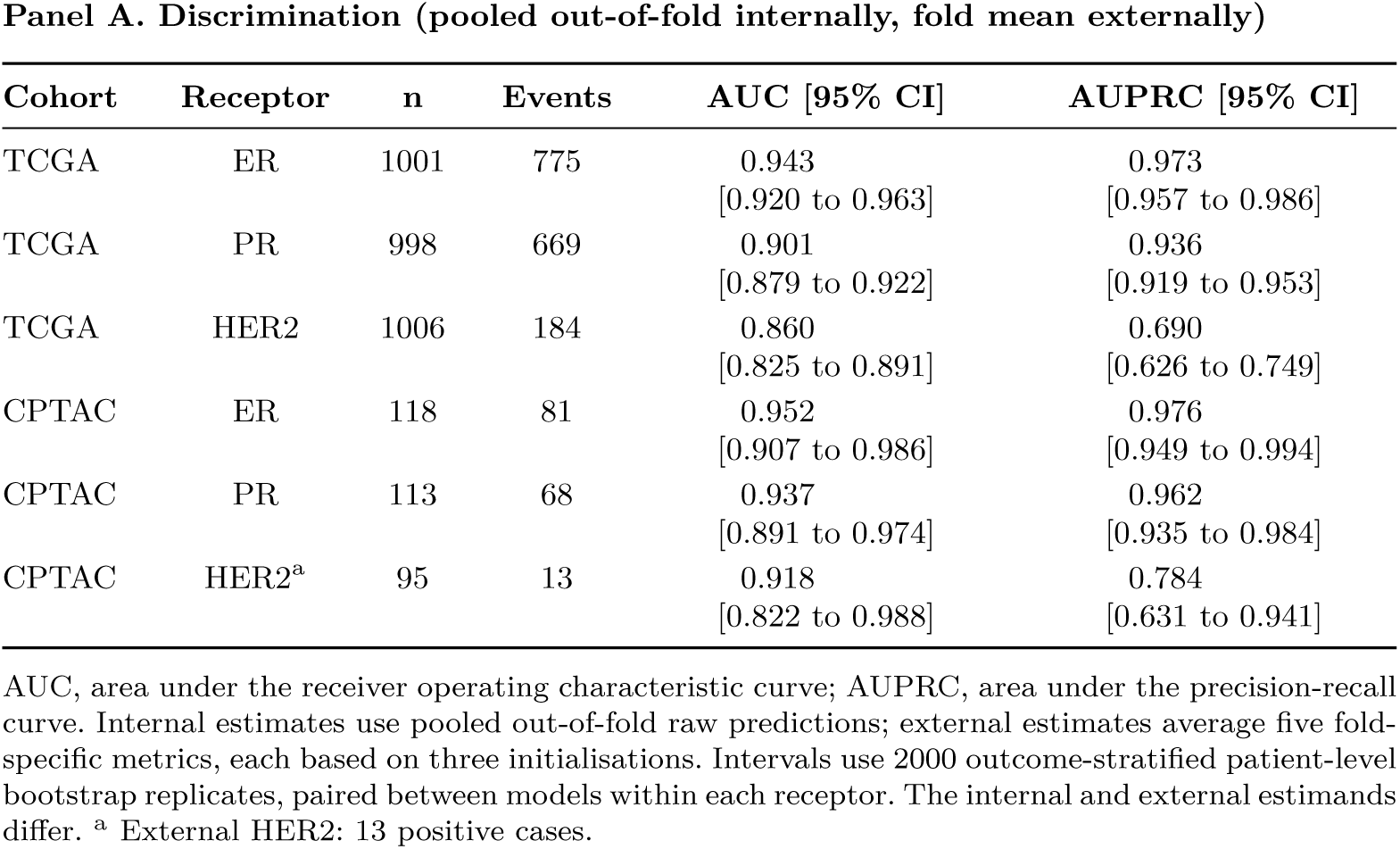
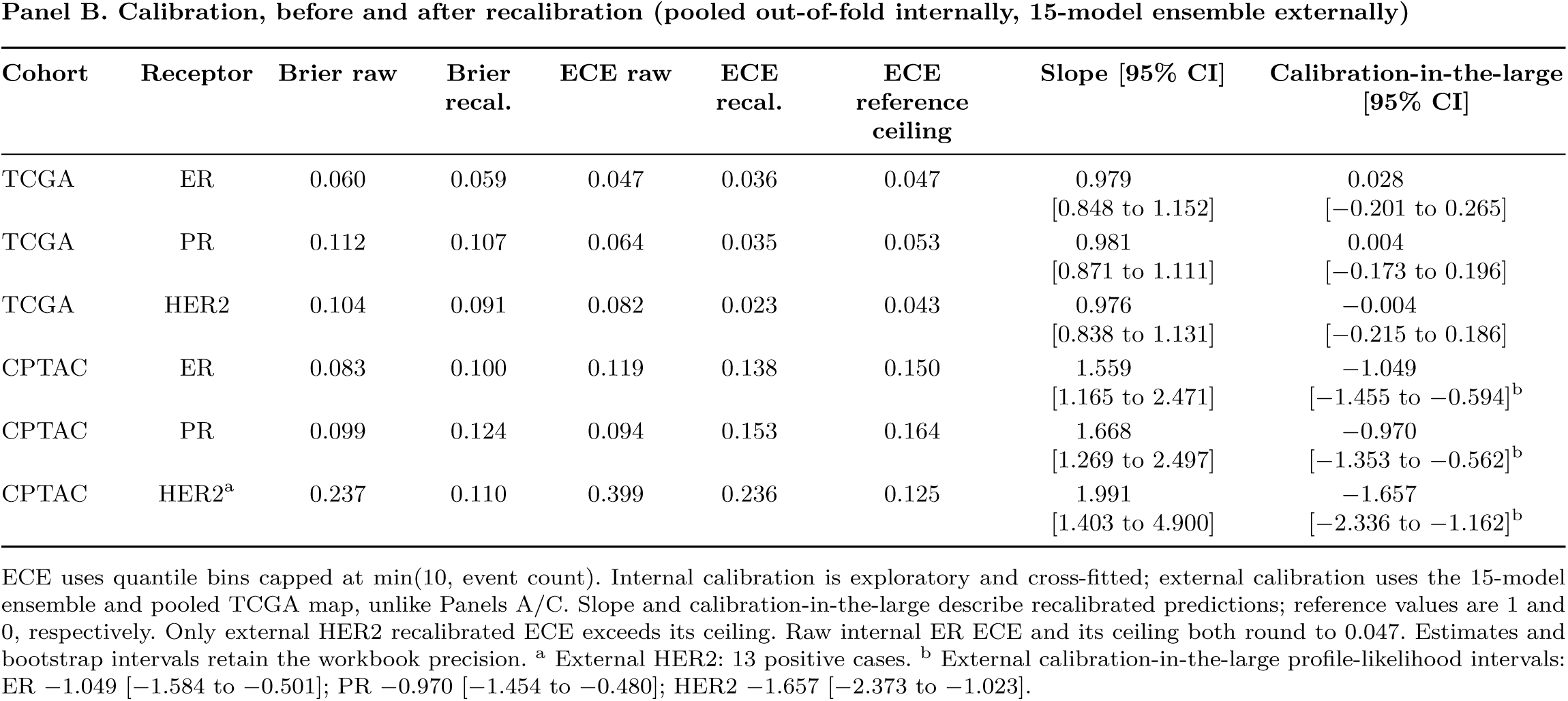
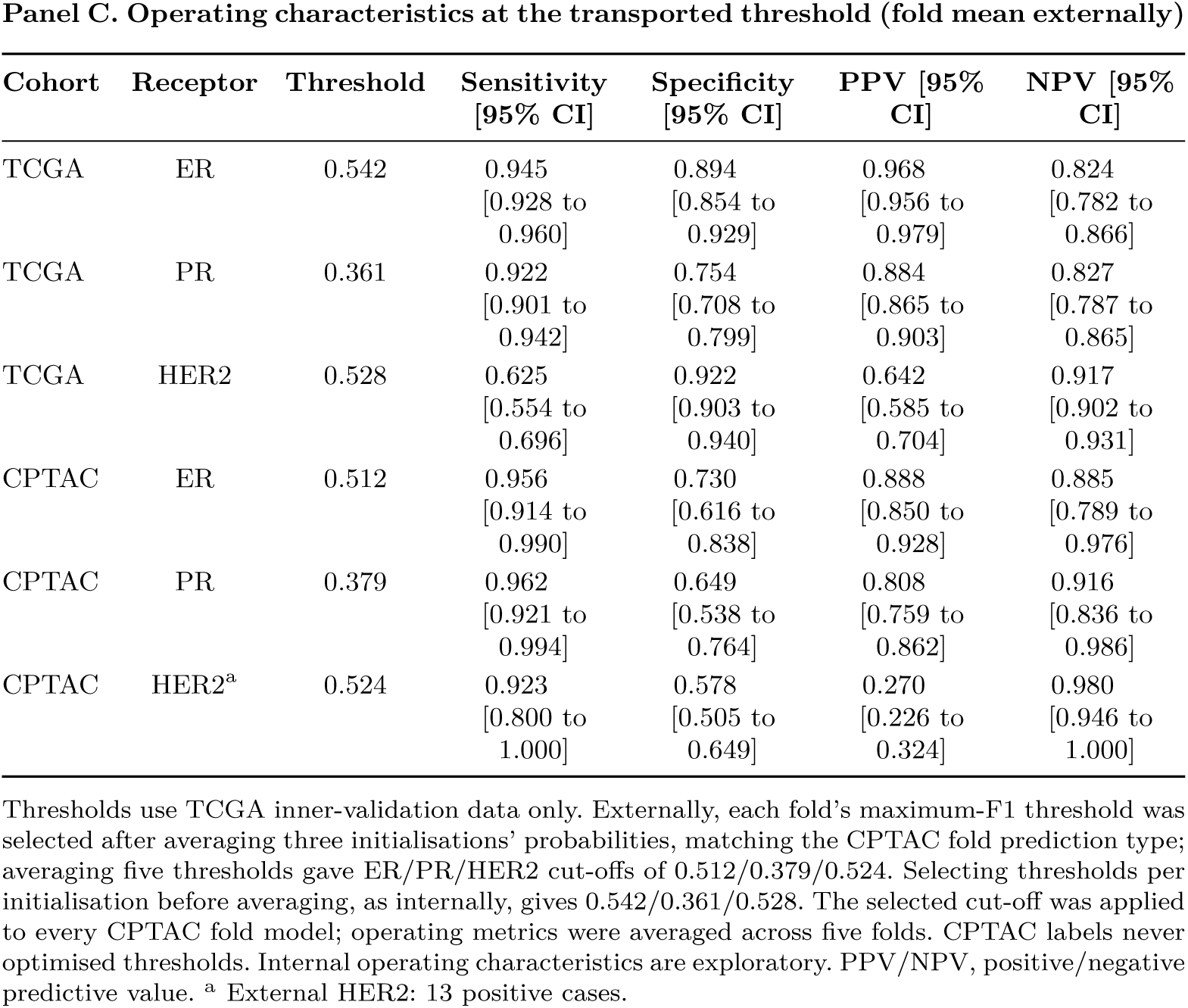
Discrimination, calibration and operating characteristics.

| Panel A. Discrimination (pooled out-of-fold internally, fold mean externally) |  |  |  |  |  |  |  |  |
|---|---|---|---|---|---|---|---|---|
| Cohort | Receptor | n | Events | AUC [95% CI] | AUPRC [95% CI] |  |  |  |
| TCGA | ER | 1001 | 775 | 0.943<br>[0.920 to 0.963] | 0.973<br>[0.957 to 0.986] |  |  |  |
| TCGA | PR | 998 | 669 | 0.901<br>[0.879 to 0.922] | 0.936<br>[0.919 to 0.953] |  |  |  |
| TCGA | HER2 | 1006 | 184 | 0.860<br>[0.825 to 0.891] | 0.690<br>[0.626 to 0.749] |  |  |  |
| CPTAC | ER | 118 | 81 | 0.952<br>[0.907 to 0.986] | 0.976<br>[0.949 to 0.994] |  |  |  |
| CPTAC | PR | 113 | 68 | 0.937<br>[0.891 to 0.974] | 0.962<br>[0.935 to 0.984] |  |  |  |
| CPTAC | HER2<sup>a</sup> | 95 | 13 | 0.918<br>[0.822 to 0.988] | 0.784<br>[0.631 to 0.941] |  |  |  |
| Panel B. Calibration, before and after recalibration (pooled out-of-fold internally, 15-model ensemble externally) |  |  |  |  |  |  |  |  |
| Cohort | Receptor | Brier raw | Brier recal. | ECE raw | ECE recal. | ECE reference ceiling | Slope [95% CI] | Calibration-in-the-large [95% CI] |
| TCGA | ER | 0.060 | 0.059 | 0.047 | 0.036 | 0.047 | 0.979<br>[0.848 to 1.152] | 0.028<br>[−0.201 to 0.265] |
| TCGA | PR | 0.112 | 0.107 | 0.064 | 0.035 | 0.053 | 0.981<br>[0.871 to 1.111] | 0.004<br>[−0.173 to 0.196] |
| TCGA | HER2 | 0.104 | 0.091 | 0.082 | 0.023 | 0.043 | 0.976<br>[0.838 to 1.131] | −0.004<br>[−0.215 to 0.186] |
| CPTAC | ER | 0.083 | 0.100 | 0.119 | 0.138 | 0.150 | 1.559<br>[1.165 to 2.471] | −1.049<br>[−1.455 to −0.594]<sup>b</sup> |
| CPTAC | PR | 0.099 | 0.124 | 0.094 | 0.153 | 0.164 | 1.668<br>[1.269 to 2.497] | −0.970<br>[−1.353 to −0.562]<sup>b</sup> |
| CPTAC | HER2<sup>a</sup> | 0.237 | 0.110 | 0.399 | 0.236 | 0.125 | 1.991<br>[1.403 to 4.900] | −1.657<br>[−2.336 to −1.162]<sup>b</sup> |
| Panel C. Operating characteristics at the transported threshold (fold mean externally) |  |  |  |  |  |  |  |  |
| Cohort | Receptor | Threshold | Sensitivity [95% CI] | Specificity [95% CI] | PPV [95% CI] | NPV [95% CI] |  |  |
| TCGA | ER | 0.542 | 0.945<br>[0.928 to 0.960] | 0.894<br>[0.854 to 0.929] | 0.968<br>[0.956 to 0.979] | 0.824<br>[0.782 to 0.866] |  |  |
| TCGA | PR | 0.361 | 0.922<br>[0.901 to 0.942] | 0.754<br>[0.708 to 0.799] | 0.884<br>[0.865 to 0.903] | 0.827<br>[0.787 to 0.865] |  |  |
| TCGA | HER2 | 0.528 | 0.625<br>[0.554 to 0.696] | 0.922<br>[0.903 to 0.940] | 0.642<br>[0.585 to 0.704] | 0.917<br>[0.902 to 0.931] |  |  |
| CPTAC | ER | 0.512 | 0.956<br>[0.914 to 0.990] | 0.730<br>[0.616 to 0.838] | 0.888<br>[0.850 to 0.928] | 0.885<br>[0.789 to 0.976] |  |  |
| CPTAC | PR | 0.379 | 0.962<br>[0.921 to 0.994] | 0.649<br>[0.538 to 0.764] | 0.808<br>[0.759 to 0.862] | 0.916<br>[0.836 to 0.986] |  |  |
| CPTAC | HER2<sup>a</sup> | 0.524 | 0.923<br>[0.800 to 1.000] | 0.578<br>[0.505 to 0.649] | 0.270<br>[0.226 to 0.324] | 0.980<br>[0.946 to 1.000] |  |  |

AUC, area under the receiver operating characteristic curve; AUPRC, area under the precision-recall curve. Internal estimates use pooled out-of-fold raw predictions; external estimates average five fold-specific metrics, each based on three initialisations. Intervals use 2000 outcome-stratified patient-level bootstrap replicates, paired between models within each receptor. The internal and external estimands differ. <sup>a</sup> External HER2: 13 positive cases.
ECE uses quantile bins capped at min(10, event count). Internal calibration is exploratory and cross-fitted; external calibration uses the 15-model ensemble and pooled TCGA map, unlike Panels A/C. Slope and calibration-in-the-large describe recalibrated predictions; reference values are 1 and 0, respectively. Only external HER2 recalibrated ECE exceeds its ceiling. Raw internal ER ECE and its ceiling both round to 0.047. Estimates and bootstrap intervals retain the workbook precision. <sup>a</sup> External HER2: 13 positive cases. <sup>b</sup> External calibration-in-the-large profile-likelihood intervals: ER −1.049 [−1.584 to −0.501]; PR −0.970 [−1.454 to −0.480]; HER2 −1.657 [−2.373 to −1.023].
Thresholds use TCGA inner-validation data only. Externally, each fold’s maximum-F1 threshold was selected after averaging three initialisations’ probabilities, matching the CPTAC fold prediction type; averaging five thresholds gave ER/PR/HER2 cut-offs of 0.512/0.379/0.524. Selecting thresholds per initialisation before averaging, as internally, gives 0.542/0.361/0.528. The selected cut-off was applied to every CPTAC fold model; operating metrics were averaged across five folds. CPTAC labels never optimised thresholds. Internal operating characteristics are exploratory. PPV/NPV, positive/negative predictive value. <sup>a</sup> External HER2: 13 positive cases.

### 3.3 Incremental value and the primary comparison

Macro-AUC was 0.901 internally and 0.936 externally for the multimodal model, compared with 0.909 and 0.957 for three-transcript logistic regression (Table 3). The external fold-mean primary difference was −0.021 [−0.046 to 0.001]. The internal pooled out-of-fold difference was −0.007 [−0.022 to 0.007], and the external 15-model ensemble difference was −0.012 [−0.038 to 0.011]. The regularised linear model using the 5000 highest-variance genes [36] had higher macro-AUC than the transcriptomic neural network in both cohorts. Image–gene attention fusion changed internal macro-AUC by 0.011 [0.001 to 0.021], compared with −0.004 [−0.012 to 0.003] externally. These component comparisons were exploratory.

**Table 3.** Modality contribution and incremental value of model components.

| Model | TCGA | CPTAC | Scope |
|---|---|---|---|
| R0 Age + stage logistic | 0.538 | 0.532 | macro |
| R1 ESR1-only logistic | 0.959 | 0.962 | ER only |
| R2 PGR-only logistic | 0.919 | 0.966 | PR only |
| R3 ERBB2-only logistic | 0.849 | 0.957 | HER2 only |
| R4 Three-transcript logistic (ESR1, PGR, ERBB2) | 0.909 | 0.957 | macro |
| R5 Elastic net, 5000 highest-variance genes | 0.906 | 0.949 | macro |
| R5b Selected-panel ridge | 0.911 | 0.937 | macro |
| R6 Transcriptomic neural network | 0.887 | 0.931 | macro |
| R7 WSI-only | 0.803 | 0.783 | macro |
| R8 Gene+WSI late fusion | 0.891 | 0.935 | macro |
| R9 Gene+WSI self-attention fusion | 0.898 | 0.928 | macro |
| R10 Multimodal model (full) | 0.901 | 0.936 | macro |
| R4b Three-transcript, focal/asymmetric loss | 0.908 | 0.957 | macro |
| S1 Multimodal model, cross-entropy | 0.904 | 0.945 | macro |
| **Paired macro-AUC differences [descriptive 95% bootstrap intervals]** |  |  |  |
| Neural versus elastic-net estimator (R6 − R5) | −0.019 [−0.030 to −0.009] | −0.017 [−0.036 to −0.002] |  |
| Wider transcriptome (R5 − R4) | −0.002 [−0.012 to +0.008] | −0.008 [−0.034 to +0.017] |  |
| WSI, via attention fusion over R6 (R9 − R6) | +0.011 [+0.001 to +0.021] | −0.004 [−0.012 to +0.003] |  |
| Attention over late fusion (R9 − R8) | +0.008 [−0.002 to +0.017] | −0.007 [−0.023 to +0.007] |  |
| Clinical (R10 − R9) | +0.003 [−0.004 to +0.010] | +0.008 [−0.0003 to +0.018] |  |
| Primary comparison: multimodal − three-transcript (R10 − R4) | −0.007 [−0.022 to +0.007] | −0.021 [−0.046 to +0.001] |  |
| Same comparison, 15-model ensemble (sensitivity) | — | −0.012 [−0.038 to +0.011] |  |

Within each cohort, all models use the same calculation method: pooled out-of-fold internally and fold mean externally, except for the explicitly labelled ensemble sensitivity row. Single-transcript rows report receptor-specific AUC rather than macro-AUC. The external fold-mean R10 − R4 contrast is primary; internal, ensemble and component comparisons are exploratory. Macro-AUC averages the three receptor-specific AUCs on their labelled analysis sets. Bootstrap indices were shared between models within each receptor, but macro replicates were combined by replicate index without joint patient resampling across outcomes. The reported 95% macro-AUC intervals are therefore descriptive and are not used to establish superiority, equivalence or non-inferiority. Differences are based on the source estimates rather than subtraction of rounded displayed values.

### 3.4 Dependence on the 17q12–q21 region

Removal of ERBB2 alone changed internal HER2 AUC by −0.004 [−0.024 to 0.018]. Removing the six-gene 17q12 core, extended 29-transcript 17q12–q21 region or full predefined label-proximal set reduced internal HER2 AUC, with receptor-specific intervals excluding zero (Table 4). In the prespecified secondary external contrast, removal of the extended 29-transcript 17q12–q21 region reduced HER2 AUC from 0.918 to 0.875 (difference −0.042 [−0.089 to −0.007]). Retaining only ESR1, PGR and ERBB2 in the transcriptomic branch of the multimodal model (G6) increased external macro-AUC by 0.021 [−0.003 to 0.047], with the interval interpreted descriptively. Feature-ranking analyses were interpreted as checks against expected receptor-associated biology rather than evidence of new mechanisms [40].

**Table 4.**
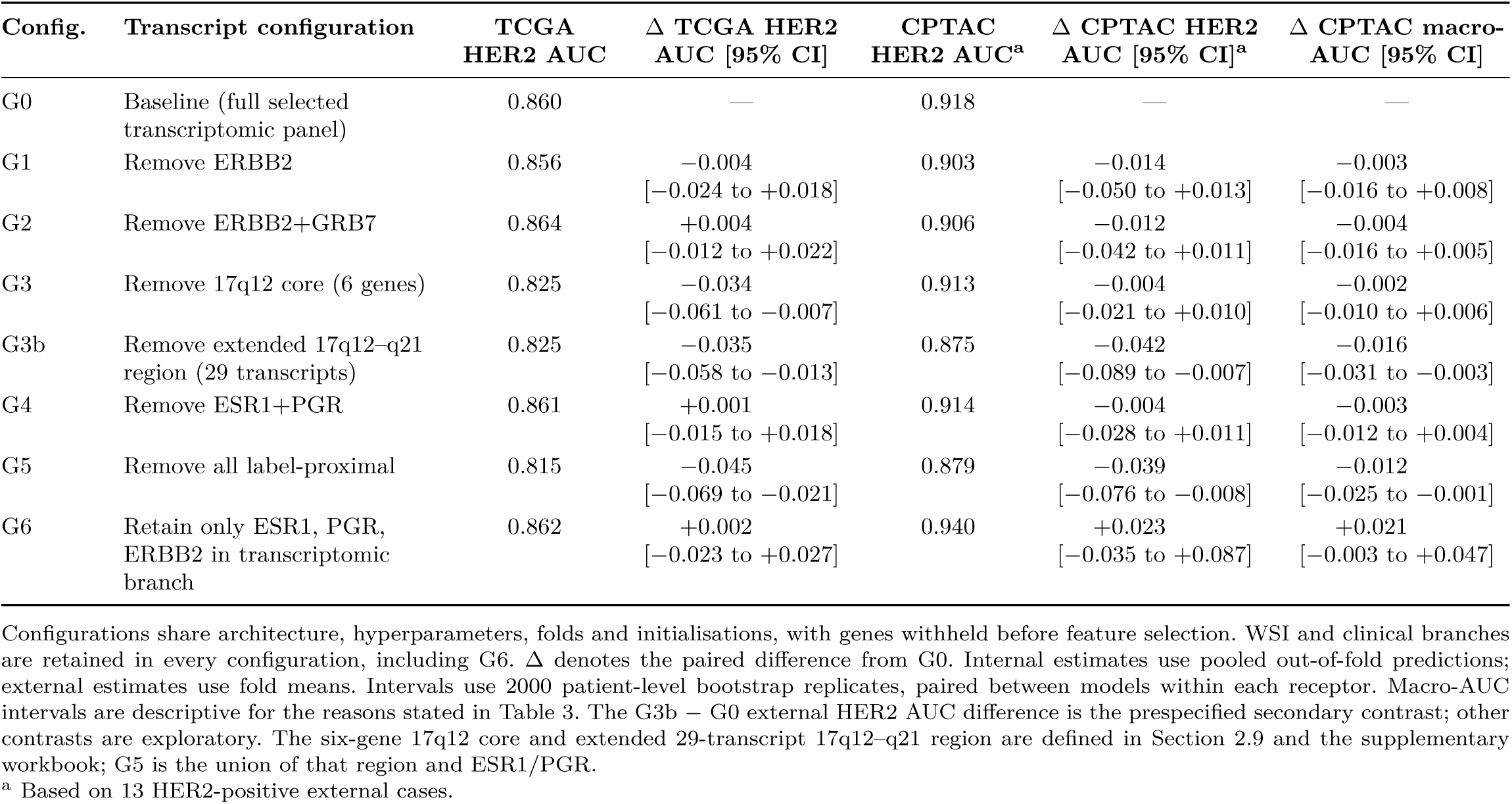
Transcript-removal and restriction experiments.

| **Config.** | **Transcript configuration** | **TCGA HER2 AUC** | **Δ TCGA HER2 AUC [95% CI]** | **CPTAC HER2 AUC<sup>a</sup>** | **Δ CPTAC HER2 AUC [95% CI]<sup>a</sup>** | **Δ CPTAC macro-AUC [95% CI]** |
|---|---|---|---|---|---|---|
| G0 | Baseline (full selected transcriptomic panel) | 0.860 | — | 0.918 | — | — |
| G1 | Remove ERBB2 | 0.856 | −0.004<br>[−0.024 to +0.018] | 0.903 | −0.014<br>[−0.050 to +0.013] | −0.003<br>[−0.016 to +0.008] |
| G2 | Remove ERBB2+GRB7 | 0.864 | +0.004<br>[−0.012 to +0.022] | 0.906 | −0.012<br>[−0.042 to +0.011] | −0.004<br>[−0.016 to +0.005] |
| G3 | Remove 17q12 core (6 genes) | 0.825 | −0.034<br>[−0.061 to −0.007] | 0.913 | −0.004<br>[−0.021 to +0.010] | −0.002<br>[−0.010 to +0.006] |
| G3b | Remove extended 17q12–q21 region (29 transcripts) | 0.825 | −0.035<br>[−0.058 to −0.013] | 0.875 | −0.042<br>[−0.089 to −0.007] | −0.016<br>[−0.031 to −0.003] |
| G4 | Remove ESR1+PGR | 0.861 | +0.001<br>[−0.015 to +0.018] | 0.914 | −0.004<br>[−0.028 to +0.011] | −0.003<br>[−0.012 to +0.004] |
| G5 | Remove all label-proximal | 0.815 | −0.045<br>[−0.069 to −0.021] | 0.879 | −0.039<br>[−0.076 to −0.008] | −0.012<br>[−0.025 to −0.001] |
| G6 | Retain only ESR1, PGR, ERBB2 in transcriptomic branch | 0.862 | +0.002<br>[−0.023 to +0.027] | 0.940 | +0.023<br>[−0.035 to +0.087] | +0.021<br>[−0.003 to +0.047] |

Configurations share architecture, hyperparameters, folds and initialisations, with genes withheld before feature selection. WSI and clinical branches are retained in every configuration, including G6. Δ denotes the paired difference from G0. Internal estimates use pooled out-of-fold predictions; external estimates use fold means. Intervals use 2000 patient-level bootstrap replicates, paired between models within each receptor. Macro-AUC intervals are descriptive for the reasons stated in Table 3. The G3b − G0 external HER2 AUC difference is the prespecified secondary contrast; other contrasts are exploratory. The six-gene 17q12 core and extended 29-transcript 17q12–q21 region are defined in Section 2.9 and the supplementary workbook; G5 is the union of that region and ESR1/PGR.
<sup>a</sup> Based on 13 HER2-positive external cases.

### 3.5 Calibration and its transport

Calibration estimates before and after recalibration are reported in Table 2B, and calibration curves are shown in Figure 3. Internal estimates used exploratory cross-fitted pooled out-of-fold predictions; external estimates used the 15-model ensemble.

**Fig 3.**
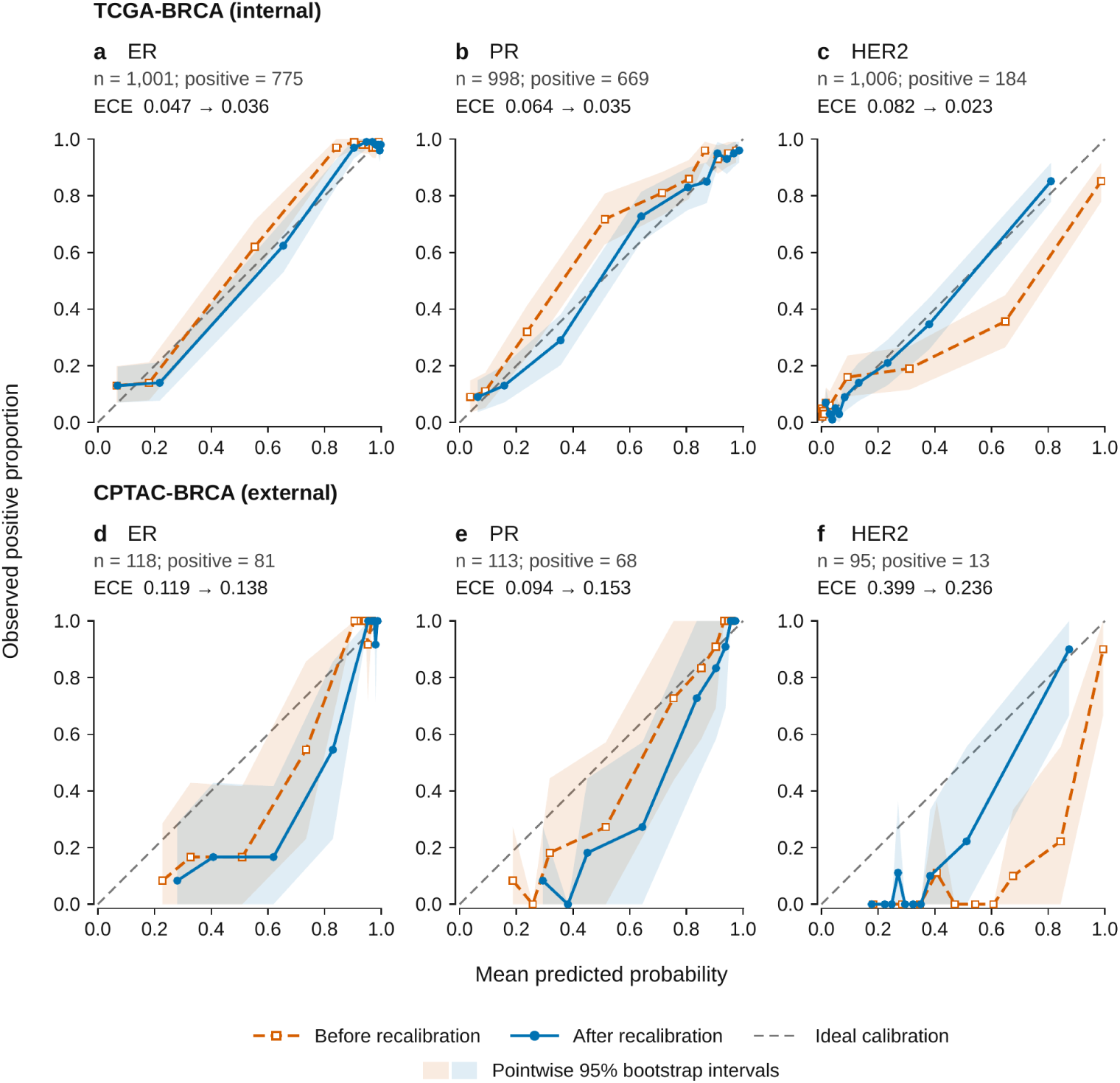
Calibration before and after Platt recalibration [37]. Each panel compares mean predicted probability with the observed receptor-positive proportion; the diagonal line indicates perfect calibration, and shaded areas show pointwise 95% bootstrap intervals. Internal panels use exploratory cross-fitted pooled out-of-fold predictions; external panels use the pooled TCGA calibration map applied to the 15-model ensemble.

### 3.6 Exploratory loss × architecture comparisons

The cross-entropy multimodal model had an external macro-AUC of 0.945, compared with 0.936 for the focal/asymmetric model [33, 34] (difference 0.010). The two three-transcript configurations both had macro-AUCs rounding to 0.957 (Table 5). With loss held constant, multimodal-minus-three-transcript differences were −0.021 under focal/asymmetric objectives and −0.011 under cross-entropy/log loss. Their difference-in-differences was −0.010.

**Table 5.**
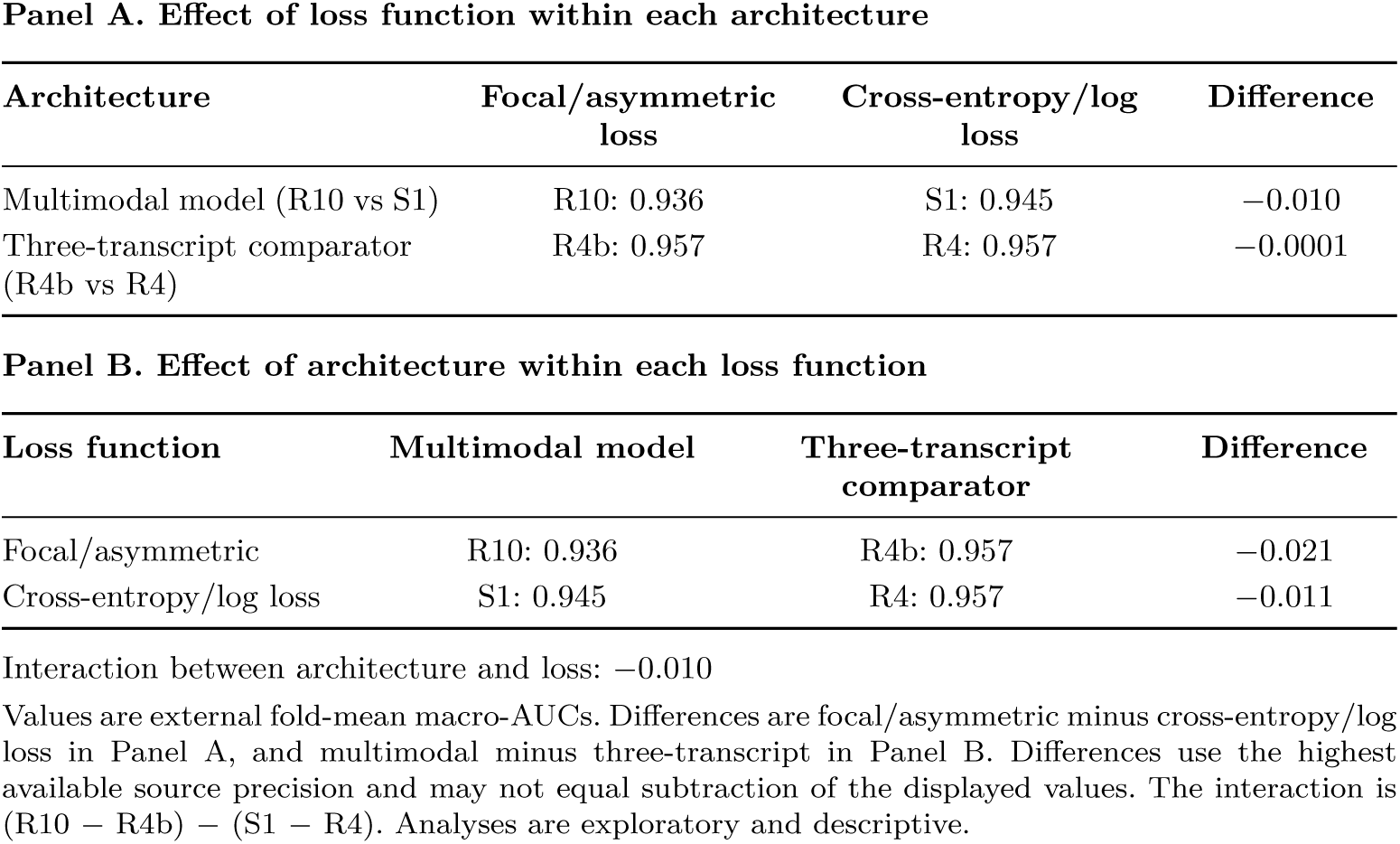
Exploratory loss × architecture comparisons of external macro-AUC.

| Panel A. Effect of loss function within each architecture |  |  |  |
|---|---|---|---|
| **Architecture** | **Focal/asymmetric loss** | **Cross-entropy/log loss** | **Difference** |
| Multimodal model (R10 vs S1) | R10: 0.936 | S1: 0.945 | −0.010 |
| Three-transcript comparator (R4b vs R4) | R4b: 0.957 | R4: 0.957 | −0.0001 |
| **Panel B. Effect of architecture within each loss function** |  |  |  |
| **Loss function** | **Multimodal model** | **Three-transcript comparator** | **Difference** |
| Focal/asymmetric | R10: 0.936 | R4b: 0.957 | −0.021 |
| Cross-entropy/log loss | S1: 0.945 | R4: 0.957 | −0.011 |

Interaction between architecture and loss: −0.010
Values are external fold-mean macro-AUCs. Differences are focal/asymmetric minus cross-entropy/log loss in Panel A, and multimodal minus three-transcript in Panel B. Differences use the highest available source precision and may not equal subtraction of the displayed values. The interaction is (R10 − R4b) − (S1 − R4). Analyses are exploratory and descriptive.

### 3.7 Head-to-head comparison with the three-transcript model

In the external fold-mean analysis, the multimodal-minus-three-transcript differences were −0.021 [−0.046 to 0.001] for macro-AUC and −0.036 [−0.100 to 0.010] for HER2 AUC (Table 6). Using the pooled TCGA calibration map, paired differences in calibration-in-the-large, ECE and Brier score were −1.519 [−1.852 to −1.213], 0.179 [0.121 to 0.223] and 0.081 [0.051 to 0.112], respectively, for HER2; corresponding ER and PR estimates are reported in Table 6. In one-initialisation-per-fold sensitivity analyses, HER2 calibration-in-the-large differences ranged from −2.380 to −1.848, and ECE differences from 0.177 to 0.241.

**Table 6.** External head-to-head comparison of the multimodal and three-transcript models.

| **Receptor / metric** | **Difference (multimodal −<br>three-transcript)<br>[95% interval]** |
|---|---|
| Macro-AUC | −0.021<br>[−0.046 to 0.001] |
| HER2 AUC<sup>a</sup> | −0.036<br>[−0.100 to 0.010] |
| ER calibration-in-the-large | −0.083<br>[−0.394 to 0.207] |
| PR calibration-in-the-large | −0.404<br>[−0.680 to −0.147] |
| HER2 calibration-in-the-large<sup>a</sup> | −1.519<br>[−1.852 to −1.213] |
| ER ECE | +0.030<br>[−0.005 to 0.065] |
| PR ECE | +0.066<br>[0.026 to 0.105] |
| HER2 ECE<sup>a</sup> | +0.179<br>[0.121 to 0.223] |
| ER Brier score | +0.027<br>[−0.001 to 0.056] |
| PR Brier score | +0.036<br>[0.012 to 0.058] |
| HER2 Brier score<sup>a</sup> | +0.081<br>[0.051 to 0.112] |

Differences are multimodal minus three-transcript logistic regression, calculated using paired external fold-mean metrics on identical patients and shared bootstrap resamples. Calibration contrasts use the pooled TCGA calibration map for each model. These are not differences between ensemble estimates in Table 2B. Intervals are based on 2000 patient-level bootstrap replicates; the macro-AUC interval is descriptive for the reasons stated in Table 3. Positive ECE and Brier-score differences indicate higher probability error for the multimodal model. <sup>a</sup> External HER2 estimates are based on 13 positive cases.

### 3.8 Decision-curve analysis (exploratory)

At a threshold of 0.200, internal HER2 net benefit was 0.104 for the multimodal model, 0.101 for the three-transcript model and 0.103 for the selected-panel ridge model, with paired-difference intervals including zero. For external 15-model ensemble predictions, the corresponding estimates were −0.063, 0.082 and 0.090. Multimodal-minus-comparator differences were −0.145 [−0.182 to −0.100] and −0.153 [−0.190 to −0.108], respectively. Treat-all net benefit was −0.021 internally and −0.079 externally (Figure 4). These exploratory comparisons were not linked to a specified clinical action.

**Fig 4.**
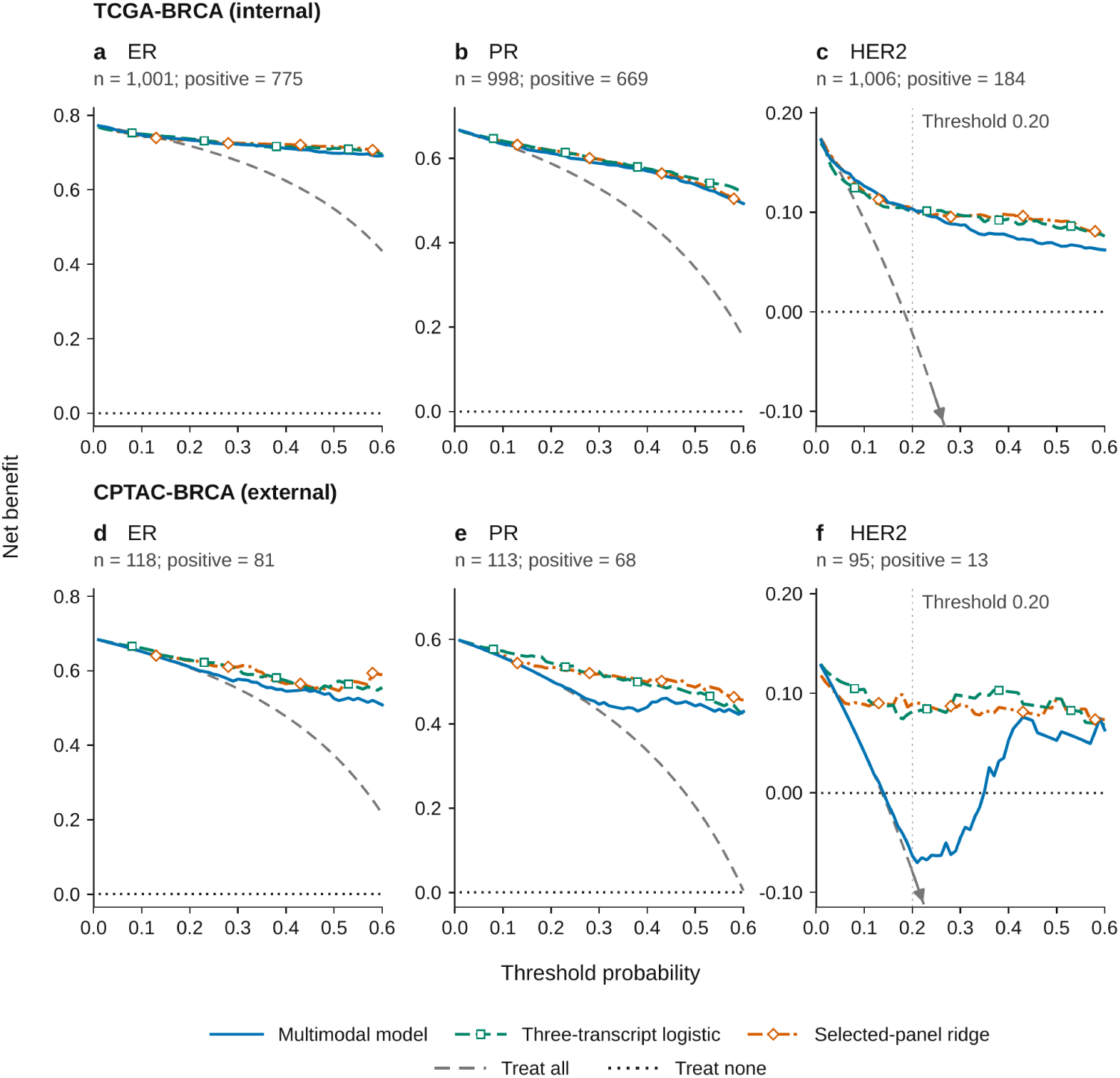
Exploratory decision-curve analysis by cohort and receptor. Internal panels use recalibrated pooled out-of-fold predictions; external panels use recalibrated ensemble predictions.

### 3.9 Reference-standard provenance and label definition

High-confidence internal HER2 labels were available for 677 patients with 135 events, compared with 329 patients and 49 events in the remaining stratum. AUCs were 0.870 and 0.828 (difference 0.043 [−0.040 to 0.131]), and AUPRCs were 0.717 and 0.623 (difference 0.094 [−0.052 to 0.237]). In the separate comparison of ISH-derived with curated-IHC labels, AUCs were 0.809 and 0.780 (difference 0.029 [−0.092 to 0.156]), and AUPRCs were 0.561 and 0.566 (difference −0.004 [−0.193 to 0.188]).

All provenance-comparison intervals included zero. AUCs under label rules L0, L1 and L2 were 0.860, 0.857 and 0.870, respectively. L1 changed 0.8% of labels, below the prespecified 2% retraining threshold. L2 used the same 677 patients and raw predictions as the high-confidence stratum and yielded identical point estimates.

### 3.10 Sensitivity analyses

Across three five-fold partitions, internal macro-AUC varied over a range of 0.006 and HER2 AUC over 0.003; corresponding exploratory initialisation ranges were 0.012 and 0.025. Training AUC exceeded held-out AUC by 0.053, 0.095 and 0.127 for ER, PR and HER2. Of 120 sensitivity-analysis units, 108 were computable and 14 changed sign or whether the interval included zero. Ten involved a normalisation variant that failed its expression-unit comparability criterion, three involved a change in aggregation method and one belonged to an exploratory analysis. These results indicate sensitivity of some estimates rather than uniform stability. The proposed single-patient normalisation variant failed the expression-unit comparability assessment and exceeded the 1% clipping criterion.

### 3.11 Validity and reproducibility

All ten programmed leakage checks passed in the internal audit. Submitting site was recoverable from TCGA image features with accuracy 0.951, compared with a majority-class baseline of 0.144 [38]. Internal replay reproduced predictions and coefficients, and re-derivation of 5400 summary values from 481 artefacts recorded no discrepancy. The audit chain contained one CPTAC scoring event after the internal freeze tag and two subsequent label-read entries for reporting. These checks document reproducibility within the reanalysis; they do not establish independence from earlier CPTAC exposure.

## 4 Discussion

### 4.1 Principal findings

The multimodal model achieved high discrimination for ER, PR and HER2 across TCGA-BRCA and CPTAC-BRCA, but did not demonstrate a clear incremental discrimination advantage over logistic regression using ESR1, PGR and ERBB2. The external macro-AUC point estimate was lower for the multimodal model, and the descriptive interval for the primary comparison included zero (Table 3); this does not establish superiority of either model, equivalence or non-inferiority. The study therefore primarily informs the incremental value of model complexity rather than identifying a superior architecture.

Regional transcript removal reduced HER2 discrimination within the evaluated pipeline, while calibration and paired probability-error comparisons revealed limitations not apparent from AUC alone. External 15-model ensemble probabilities overpredicted all three receptors, and paired fold-mean ECE and Brier-score comparisons favoured the three-transcript model for HER2 and PR (Tables 2B and 6). These results concern predictive performance; the proposed laboratory quality-assurance workflow was not directly evaluated.

### 4.2 Incremental value and molecular dependence

Previous studies have shown that H&E morphology contains information associated with receptor status [6–11], while transcriptomic markers such as ESR1, PGR and ERBB2 provide strong molecular signals [14, 15]. In our comparisons, transcript-based models generally discriminated better than the image-only model. The relevant question was therefore whether adding histology, clinical variables and nonlinear multimodal modelling provided value beyond an informative molecular baseline, particularly in settings where tumour RNA measurements are already available.

Within this study, the modest internal gain from image–gene attention fusion was not reproduced externally. A regularised linear model using the 5,000 highest-variance genes also achieved higher macro-AUC than the transcriptomic neural network in both cohorts, consistent with previous evidence that greater model complexity does not necessarily improve clinical prediction performance [21]. Restricting only the transcriptomic branch of the multimodal network to ESR1, PGR and ERBB2, while retaining WSI and clinical inputs, did not show superiority over the full selected transcriptomic panel because the descriptive interval for the external difference included zero (Table 4). Overall, these comparisons provide no consistent evidence that a broader transcriptomic panel or greater neural-network complexity improved discrimination in this setting. Changing the training loss produced the same general pattern: the multimodal model remained numerically below the three-transcript comparator under both loss functions, while the difference between these effects was descriptive rather than evidence of a confirmed interaction (Table 5).

The transcript-removal experiments nevertheless clarified where predictive information may reside. Removing ERBB2 alone had little observed effect on internal HER2 AUC, whereas removing the extended 29-transcript 17q12–q21 region reduced HER2 discrimination in both cohorts (Table 4). This pattern is compatible with predictive information distributed across ERBB2 and neighbouring correlated transcripts, as described in genomic studies of HER2-positive breast cancer [16–18]. Feature rankings were treated as checks against established receptor-associated biology rather than evidence of a newly identified mechanism [40]. These removal and restriction analyses answer different questions and do not establish a causal role for individual genes; residual discrimination may reflect correlated transcription, tumour composition, morphology or cohort structure.

### 4.3 Calibration and probability performance

High discrimination did not imply stable probability estimates across cohorts. Calibration concerns agreement between predicted probabilities and observed outcomes [22], and the TCGA-derived recalibration mapping did not eliminate external overprediction. For the external 15-model ensemble, calibration-in-the-large was negative for all three receptors and calibration slopes remained above one (Table 2B), indicating that the probability scale did not transport intact despite preserved discrimination. HER2 recalibrated ECE exceeded its ECE reference ceiling, whereas ER and PR ECE did not; however, the latter should not be interpreted as well calibrated because their slope and calibration-in-the-large estimates still showed discrepancies. The ECE ceiling is a finite-sample reference under specified simulation scenarios rather than a universal calibration test.

Recalibration also had heterogeneous effects: external HER2 Brier score improved, whereas ER and PR Brier scores increased (Table 2B; Figure 3). Possible contributors include differences in prevalence, expression platforms, case mix, historical reference labels and cohort-level normalisation [41, 42], but the present analyses cannot separate these effects from those of the fitted models or training objectives. Internal pooled out-of-fold estimates, external fold-mean metrics and external ensemble calibration summarise different prediction objects and should therefore not be treated as interchangeable.

The paired comparisons in Table 6 reinforce this distinction. For HER2, the AUC-difference interval included zero, while ECE and Brier-score differences favoured the three-transcript model; the calibration-in-the-large contrast was also consistent with greater overprediction by the multimodal model. PR probability-error comparisons similarly favoured the three-transcript model, whereas ER intervals included zero. These findings do not imply perfect calibration of the comparator, nor do they establish superiority in discrimination. The one-initialisation-per-fold HER2 sensitivity analysis retained the direction of the calibration differences, but this remains a sensitivity analysis within the same cohort rather than independent confirmation.

### 4.4 Clinical implications and next steps

Exploratory decision-curve analysis illustrated that the consequences of predicted probabilities depend on the chosen threshold [23]. At a threshold of 0.200, external 15-model ensemble HER2 net benefit for the multimodal model was below that of the three-transcript and selected-panel ridge comparators. This result is specific to the evaluated cohort, threshold and prediction convention and should not be interpreted as evidence of clinical harm. No clinical action, review cost or patient outcome was attached to any threshold, so clinical utility was not established.

Laboratory quality assurance therefore remains a hypothetical application in settings already generating digital pathology and tumour RNA sequencing. The transported-threshold results classify receptor positivity rather than detect erroneous laboratory reports, and external HER2 estimates were based on only 13 positive cases (Table 2C). Future work should evaluate a fully specified pipeline in a previously unobserved, multi-institutional cohort with contemporary adjudicated labels, more HER2-positive cases, deployment-compatible expression preprocessing and strong molecular comparators. The laboratory quality-assurance workflow should be tested prospectively through an explicit discordance-review pathway, with IHC/ISH retained as the reference assessment [2, 3, 7], locally assessed calibration and clinically justified thresholds.

### 4.5 Strengths and limitations

Strengths include patient-level evaluation, matched comparator analyses with paired predictions, and separate assessment of discrimination, calibration and exploratory decision curves. Label-provenance analyses, sensitivity checks and internal replay further improve transparency of the reanalysis [24, 25].

Several limitations constrain interpretation. CPTAC was small, with only 13 HER2-positive cases, and had been examined during earlier development; it therefore provides cross-cohort evaluation rather than fully independent validation. Historical HER2 annotations were heterogeneous, the binary outcome did not distinguish lower-expression categories [3, 5], and stable subgroup or fairness analyses were not feasible. Alternative HER2 label rules were evaluated without retraining, so they assess sensitivity of the reported metrics rather than the full development process. Repeated five-fold partitions under fixed configurations did not reproduce complete model selection within outer folds, and training-test AUC differences should not be interpreted as estimates of selection-induced optimism. Internal recalibration reused a common cross-validation partition and should therefore remain exploratory. Macro-AUC bootstrap intervals were descriptive because receptor-specific replicates were combined without joint patient resampling, and predictive-value intervals were conditional on the observed prevalence. Alternative resampling sensitivity was available for only part of the analysis.

Finally, cohort-level rank transformation limits isolated single-patient deployment, and the reference-based alternative failed its expression-unit comparability criterion. Some sensitivity estimates changed direction or whether intervals included zero, and recoverable image-site signatures remain a potential source of non-biological signal [38]. The clinical branch was restricted to age and stage. Retained panels, predictions and replay records support reproducibility of the reported implementation, but do not substitute for independent reproduction or prospective evaluation.

## 5 Conclusions

This study addressed three linked objectives. First, the multimodal model achieved high discrimination in both cohorts but did not demonstrate clear incremental pre-dictive value over a parsimonious logistic model using ESR1, PGR, and ERBB2; added image, clinical, and nonlinear modelling components did not yield a consistent transportable advantage. Second, transcript-removal experiments showed that HER2 performance depended in part on information within the extended 29-transcript 17q12–q21 region, although these analyses do not establish a biological mechanism or causal contribution of individual genes. Third, predicted probabilities did not transport consistently across cohorts: external calibration remained problematic, particularly for HER2, and paired probability-error comparisons favoured the three-transcript model for HER2 and PR. Exploratory decision-curve findings were threshold dependent and did not establish clinical utility. These results support benchmarking complex multimodal models against informative molecular baselines and evaluating calibration alongside discrimination. Prior CPTAC exposure, limited external HER2 events, and deployment-related preprocessing constraints warrant independent evaluation before any proposed laboratory quality-assurance workflow is considered for clinical use.

## List of abbreviations

AJCC: American Joint Committee on Cancer
AUC: area under the receiver operating characteristic curve
AUPRC: area under the precision-recall curve
CI: confidence interval
CIL: calibration-in-the-large
CLS: classification token
CPM: counts per million
CPTAC-BRCA: Clinical Proteomic Tumor Analysis Consortium breast cancer cohort
CV: cross-validation
DCA: decision-curve analysis
ECE: expected calibration error
ER: estrogen receptor
FISH: fluorescence in situ hybridisation
GDC: Genomic Data Commons
GELU: Gaussian error linear unit
H&E: haematoxylin and eosin
HER2: human epidermal growth factor receptor 2
IHC: immunohistochemistry
IQR: interquartile range
ISH: in situ hybridisation
LN: layer normalisation
MHA: multi-head attention
MLP: multilayer perceptron
NPV: negative predictive value
OOF: out-of-fold
PPV: positive predictive value
PR: progesterone receptor
ReLU: rectified linear unit
RNA-seq: RNA sequencing
TCGA-BRCA: The Cancer Genome Atlas breast cancer cohort
WSI: whole-slide image.

## Supplementary material

Supplementary results workbook: BMC_MIDM_rebuild_results_summary_with_defi nitions.xlsx. The workbook contains definitions, gene sets, cohort and label provenance, comparator and transcript-removal results, calibration and operating characteristics, paired contrasts, decision-curve summaries, sensitivity analyses and reproducibility records.

## Declarations

### Ethics approval and consent to participate

This study was a secondary analysis of de-identified TCGA-BRCA and CPTAC-BRCA repository data. No participants were recruited or contacted, no new specimens were collected, and no intervention was undertaken. Ethics approval and additional informed consent were not required for this secondary analysis. Data were accessed and analysed in accordance with the applicable TCGA/CPTAC data-access and data-use policies. No attempt was made to re-identify any participant.

### Consent for publication

Not applicable. No individually identifiable participant data, images or videos are presented.

### Availability of data and materials

TCGA-BRCA clinical, RNA-sequencing and diagnostic-slide data were obtained from the Genomic Data Commons, and CPTAC-BRCA data from the Proteomic Data Commons. The study generated model and analysis outputs but no new biological specimens. The analysis code and deposited analysis records are available from the project repository [39]; the associated archive DOI is https://doi.org/10.5281/zenodo.22711622. The repository includes code, documentation, tables and figures under the MIT licence. Original input data remain subject to their source repositories’ access and use conditions. Derived per-patient feature bags and trained checkpoints are not redistributed in the public repository.

### Authors’ contributions

BMT, VAN: Conceptualization; Methodology; Software; Formal analysis.

QAD, NMHN, QAN, HNT: Conceptualization; Formal analysis; Writing – original draft; Writing – review & editing.

DHT, TBVN: Supervision; Methodology; Writing – review & editing. All authors read and approved the final manuscript.

## Acknowledgements

The results are based in whole or in part on data generated by the TCGA Research Network and the Clinical Proteomic Tumor Analysis Consortium. The authors thank the patients and investigators who contributed to these resources.

## Additional file descriptions

**Additional file 1.** Additional_file_1_TRIPOD_AI_completed.pdf (PDF). *Completed TRIPOD+AI reporting checklist.* Official checklist with manuscript page references, item-level evidence, reporting gaps and an abstract-checklist assessment.

**Additional file 2.** Additional_file_2_PROBAST_AI_completed.pdf (PDF). *Completed PROBAST+AI self-assessment.* Outcome-specific assessments for ER, PR and HER2, distinguishing development, internal evaluation and CPTAC cross-cohort evaluation, with supporting rationales.

**Additional file 3.** BMC_MIDM_rebuild_results_summary_with_definitions.xlsx (XLSX). *Supplementary results workbook.* Definitions, gene sets, cohort and label provenance, model comparisons, calibration, operating characteristics, decision-curve summaries, sensitivity analyses and reproducibility records.

